# Correlates of head-fixed orienting movements in mouse superior colliculus and substantia nigra *pars reticulata*

**DOI:** 10.1101/2025.05.02.651955

**Authors:** Ted K. Doykos, Taylor Yamauchi, Anna Buteau, Spencer Hanson, Joshua T. Dudman, Gidon Felsen, Elizabeth A. Stubblefield

## Abstract

Orienting movements are a critical component of the natural behavioral repertoire, but their underlying neural bases are not well understood. The deep superior colliculus (dSC) integrates input from several brain regions that influence the selection of targets for orienting movements and coordinates activity among brainstem motor nuclei to initiate and execute movement. Evidence suggests that one prominent dSC input, the substantia nigra *pars reticulata* (SNr), permits movement by disinhibiting its targets, but much is unknown about the relationship between SNr activity, dSC activity, and movement. Building on increasing application of the head-fixed mouse model to elucidate the neural basis of behavior, we examined neural activity recorded in dSC and SNr with high-density probes in mice performing several variants of a sensorimotor orienting task, from our labs and in data sets curated by the International Brain Laboratory. We found that dSC and SNr were active preceding and throughout movement, across task variants, suggesting that they were engaged by the required movements. Prior to movement, SNr activity reflected the outcome of the previous trial, consistent with a role in biasing movements towards the highest value target. However, the dependence of dSC activity on movement direction was weaker than in other directional orienting behaviors, and we found little evidence for strong suppression of dSC by SNr. These results complement and extend previous findings from other orienting tasks and suggest diverse roles for modulatory input from SNr to dSC in shaping motor behavior.

## INTRODUCTION

The neural circuits responsible for selecting, initiating and executing movements have been studied for decades but remain poorly understood. Orienting movements are a fundamental component of natural behavior and provide convenient behavioral readouts of decision making in animal model studies. While many brain regions, including cortical, cerebellar and midbrain structures, have been implicated in orienting movements (Munoz & Everling, 2004), a key node in the network is the intermediate and deep layers of the superior colliculus (dSC), which receives a wide range of cell-type-specific input (Benavidez et al., 2021; Doykos et al., 2020; Sparks & Hartwich-Young, 1989), performs computations to select spatial targets for movement, and coordinates downstream motor circuits to initiate and execute movements (Basso et al., 2021; Gandhi & Katnani, 2011; Isa et al., 2021). The dSC thus provides an excellent model for understanding how integrating input from multiple systems subserves a well-defined function (Wolf et al., 2015). In particular, examining neural activity in dSC and its upstream inputs during directional orienting behavior can elucidate whether, when, and how the strength of input from specific regions depends on behaviorally-relevant variables.

A prominent inhibitory input to the dSC from the substantia nigra *pars reticulata* (SNr), an output nucleus of the basal ganglia, is proposed to permit orienting via phasic release of dSC from tonic inhibition (Deniau et al., 2007; Frost-Nylén et al., 2024)(Hikosaka & Wurtz, 1983; Rossi et al., 2016) and to modulate dSC output based on the value and kinematics of the movement (Basso & Wurtz, 1997; Hikosaka et al., 2006; Yttri & Dudman, 2018). The preponderance of this foundational work was performed in primates making saccades while either dSC or SNr activity was recorded (Basso & Sommer, 2011; Basso & Wurtz, 2002; Glimcher & Sparks, 1992; Handel & Glimcher, 1999; Hikosaka & Wurtz, 1983; Horwitz & Newsome, 2001; Wurtz & Goldberg, 1971). However, the extent to which these findings translate to other common forms of orienting movements, or to other species, is unclear. Thus, much remains unknown about how SNr influences the dSC computations underlying directional motor output.

To address this knowledge gap, we developed several variants of a head-fixed behavioral task requiring mice to perform an orienting directional forelimb movement to either the left or right. This task leverages the amenability of the mouse nervous system to cell-type-specific recording and perturbation during behavior (Luo et al., 2018), complementing work in primate and other model systems (Carandini & Churchland, 2013). We examined how SNr and dSC activity, recorded with high-density electrode arrays, related to behavioral events. In addition, we replicated our analyses on publicly available data recorded in mouse SNr and dSC during a similar behavioral task developed by the International Brain Laboratory (IBL) (International Brain Laboratory et al., 2024; The International Brain Laboratory et al., 2021). Across data sets we found that mice exhibited reward-maximizing behavioral strategies, SNr and dSC activity reflected several task-related variables throughout the trial, and that SNr modulated, rather than strongly suppressed, task-relevant dSC activity.

## METHODS

### Animals

Experimental animals consisted of 7 adult male mice: 3 transgenic VGAT mice (B6.Cg-Tg(Slc32a1-COP4*H134R/EYFP)8Gfng/J, https://www.jax.org/strain/014548), 3 Gad2Cre mice (Gad2-IRES-Cre, https://www.jax.org/strain/010802) and 1 wild-type C57/BL6 mouse (https://www.jax.org/strain/000664). Mouse lines expressing Cre-recombinase were produced by the GENSAT project (GENSAT project, Rockefeller University, New York, USA) and obtained from the MMRC (https://www.mmrrc.org). All animals were handled in accordance with guidelines approved by the Institutional Animal Care and Use Committee (IACUC) of the Janelia Research Campus or the University of Colorado.

Mice were individually housed in a humidity- and temperature-controlled room and maintained on a reversed 12-h light/dark cycle. Mice were implanted with a headcap under aseptic conditions, and all elements and remaining skull were covered with dental acrylic as described previously (Osborne & Dudman, 2014). Following one week of surgical recovery, mice received water restriction of 1.5 ml per day. Mice underwent daily health checks, and water restriction was eased if mice fell below 70% of their body weight at any time during deprivation. Mice were acclimated to head fixation and trained to lick drops of water sweetened with saccharin.

### Behavioral task

For our primary behavioral task, water-restricted mice were trained to rotate a custom-made, bi-directional wheel to the left or right with their forepaws to bring an eccentric visual stimulus to the center of their visual field in closed-loop to obtain water reward. The wheel was positioned under the forepaws of the mouse at an angle adjusted for the purpose of generating smooth, ballistic movements toward either the left or right side of the body, resulting in an orienting movement. The visual stimulus consisted of a bright, vertical bar on a black background of a video monitor centered 25-30 cm in front of the mouse. The stimulus was presented to either the left or right visual hemifield, randomly selected on each trial, at approximately 30 degrees horizontally from the nose. Choices were considered correct and were immediately rewarded from a front-mounted water port when the leading edge of the visual stimulus crossed the center point of the screen. Following correct trials, the inter-trial-interval (ITI) was 3.5 sec. Trials in which the mouse displaced the stimulus by the required magnitude but in the incorrect direction were not rewarded, and triggered an ITI of 7 sec. Training occurred once per day, and mice typically performed up to 350 to 400 trials per session. Recordings were conducted once mice reached a proficiency of performing 75% of trials correctly.

In the course of task optimization, the coupling between wheel and stimulus movement direction was opposite to that described above in 2 mice. The data collected from these mice, as well as across strains, did not meaningfully differ, and therefore data are combined across couplings and strains.

Separate groups of mice were trained on one of two variants of this primary task: 1) a blocked variant, in which the side of stimulus presentation was determined by a Markov process with a low transition rate (i.e., stimuli were presented at the same side on nearly every trial of the block for up to 35-50 trials); and 2) a movement cessation variant, in which mice were required to stop their movement on target to receive reward.

### Electrophysiological recordings

After reaching sufficient proficiency at the behavioral task (11-146 sessions), acute electrophysiological recordings were conducted. In one subset of mice, both the dSC and ipsilateral SNr were simultaneously targeted using acute Neuropixels (Imec, “option 3”) recording probes. Recordings were collected from left or right hemispheres during separate behavioral sessions. Neural signals were acquired using the Whisper recording system (Janelia Research Campus) with simultaneous voltage sampling from the wheel and digital signals from the behavior control system at 10 kHz. The neural recording data were processed and spike sorted using Kilosort and phy. In subsets of mice, either the SNr was targeted using acute “Hires” 32 channel silicone probes (APIG, Janelia Research Campus) or the dSC was targeted using 32-channel Neuronexus probes. Continuous data (0.1 Hz-7.5 kHz) were recorded with simultaneous sampling of voltage from the wheel and digital signals from the behavior control system (30 kHz sample rate, CerePlex Direct digital acquisition system, Blackrock Microsystems, Salt Lake City, Utah). Continuous voltage signals were high-pass filtered (0.5–7 kHz) offline, and events that exceeded four times the standard deviation of the continuous voltage signal were extracted (spikes). Spike sorting for individual units was performed in MATLAB using custom-written software or using Blackrock Online Spike Sorting software. Spikes were isolated according to waveform amplitude distributions and principal components of the amplitude array across the four electrodes (∼25 μm spacing) of each shank (*n* = 4) of the silicon probe array. The event times for each single-unit were aligned to movement start as extracted from the continuous voltage signal from the wheel. At the completion of neural recording experiments in each mouse, recording locations were confirmed histologically.

### Data Analysis

#### Direction preference

To examine the relationship between neural activity and movement direction, we performed a receiver operating characteristic (ROC)-based analysis that quantifies the extent to which an ideal observer could classify whether spike rate in a particular epoch corresponded to left or right choices (Crapse & Basso, 2015; Green & Swets, 1966). We defined direction preference as

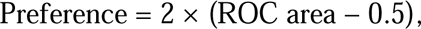

with values closer to −1 or 1 indicating a strong preference for ipsilateral or contralateral choices, respectively. A minimum of 20 trials per side was required to run the ROC analysis. Statistical significance was determined through permutation testing. Firing rates from each condition were shuffled and randomly reassigned 5,000 times, generating a distribution of preference values under the null hypothesis. The p-value was then calculated by assessing where the observed preference fell within this null distribution, reflecting the probability of observing such a preference by chance.

#### Regression Analysis

We used linear regression models to characterize the relationship between neural activity and task-related variables. For each unit, we modeled firing rates in specific epochs using reaction time (RT), choice, previous choice, and previous outcome as predictors:

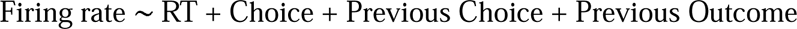

Given the potential for multicollinearity between previous choice and previous outcome, we computed the variance inflation factor (VIF) for each predictor to confirm low multicollinearity risk, which allowed us to retain both predictors in our initial model.

To determine the best-fitting model for each epoch, we calculated the Bayesian Information Criterion (BIC) for both the full model and for reduced models, each omitting one predictor. The model with RT, choice, and previous outcome emerged as providing the best fit:

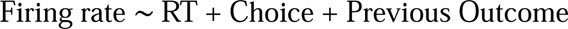

We further examined this model by testing all possible two-way interactive terms. The inclusion of interactive terms did not improve model performance, as evidenced by higher BIC values, indicating a poorer tradeoff between goodness-of-fit and model complexity.

#### Optogenetic tagging

A subset of GAD2-Cre mice was injected with FLEX-ChR2 (pAAV-Ef1a-DIO-hChR2(H134R)-EYFP) into one hemisphere of the SNr (3.5mm posterior and 1.25 mm later from bregma, 4.5 mm from the brain surface) and followed the recovery protocol described above. Prior to the behavioral session, these mice underwent optogenetic stimulation for the purpose of antidromically and optically “tagging” SNr units projecting to the ipsilateral SC. Mice were head-fixed and not permitted to begin the task until after at least one minute of photostimulation was completed. Optogenetic photostimulation was also performed for at least one minute upon completion of the behavioral task to ensure that previously tagged units were stably recorded throughout the behavioral session. While both ipsi- and contralaterally-projecting SNr-dSC neurons were assessed during recording sessions, only the ipsilaterally-projecting SNr-dSC units exhibited tagged activity, and thus functional connectivity. A laser emitting 473 nm light at a power of 4-6 mW was positioned at the surface of the brain to stimulate ChR2-expressing SNr axon terminals within the dSC and pulsed at 1 Hz, 20 ms on/980 ms off. This duty cycle was empirically determined to optimally stimulate ChR2-expressing SNr neurons (Brown et al., 2014). Recordings were obtained from the SNr during initial photostimulation, during the behavioral task, and during post-task photostimulation. By off-line spike sorting (described above), isolating single-unit events, and examining the waveform shapes of the light-elicited spikes, we determined which SNr units recorded during the behavioral session were also light-driven (i.e., expressed ChR2), and were thus putatively dSC-projecting SNr (“tagged”) units (Cardin et al., 2010; Cohen et al., 2012; Essig et al., 2021; Lima et al., 2009). For each unit, we calculated cross-correlations (MATLAB) of the average waveforms for light-driven spikes and spontaneous spikes to ensure light-driven spikes were not contaminated by noise induced by optical heating or illumination. A cross-correlation value ≥0.96 was considered a tagged unit.

## RESULTS

Mice were trained to perform a head-fixed sensorimotor task requiring the translation of a visual stimulus (bright, vertical bar) randomly displayed on the left or right periphery of a screen toward a central target by turning a rotary wheel with an orienting forepaw movement (Fig. 1a). Mice typically achieved a performance of ≥75% correct within 25-50 sessions, consistent with data collected during a similar head-fixed sensorimotor task by the International Brain Laboratory (International Brain Laboratory et al., 2024). Despite stereotyped choice performance once mice were fully trained (Fig. 1D), we observed a bimodal distribution of reaction times across trials (defined as the duration from stimulus onset to movement onset; Fig. 1E), with the shortest reaction times less than 100 ms and longer reaction times approximating 350 ms. Based on predictions of a classical speed-accuracy tradeoff framework, previously observed in similar behaviors (Luce, 1986; Rinberg et al., 2006; Shadlen & Newsome, 2001), we hypothesized that these divergent reaction times correlated with task performance.

**Figure 1.**
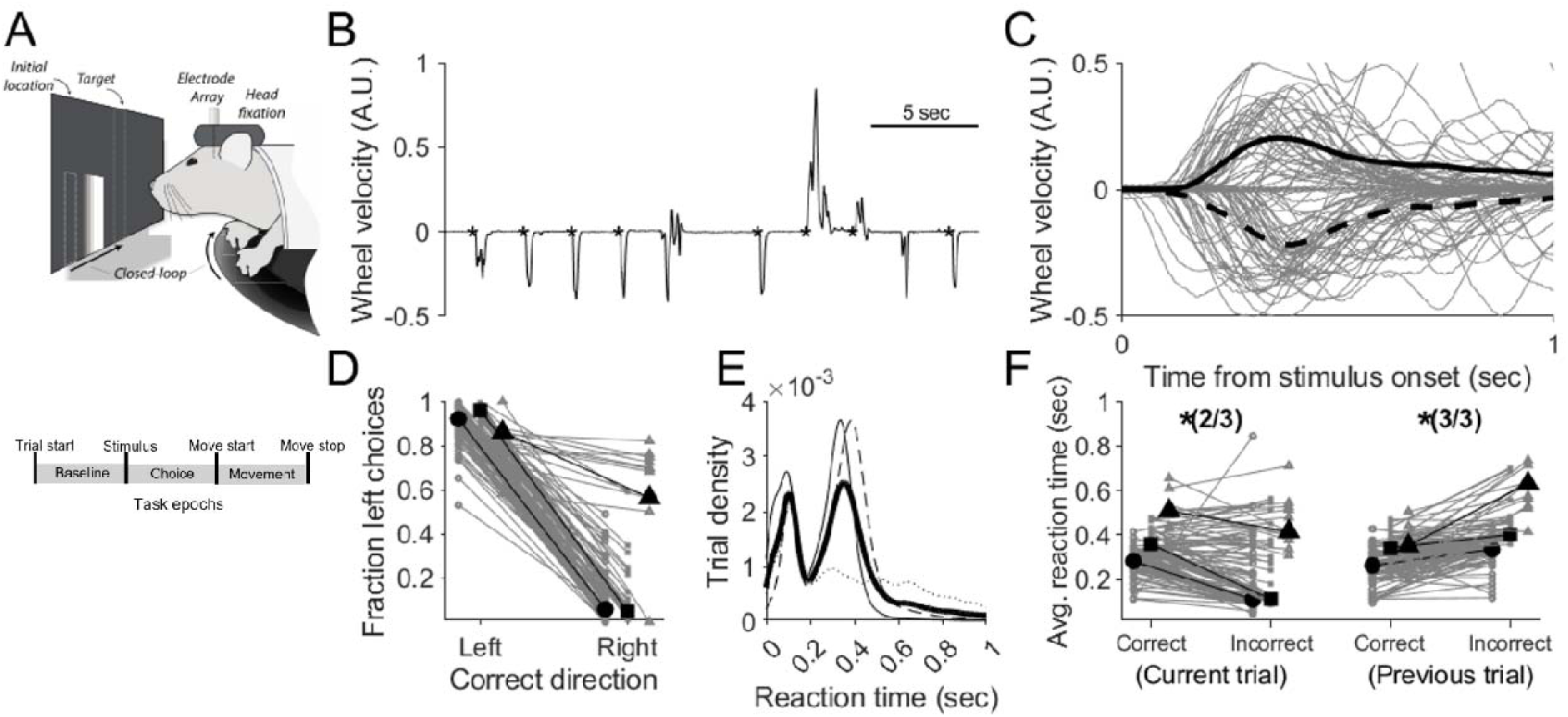
Head-fixed mice perform a forelimb orienting task with high accuracy and variable reaction times. A) Behavioral setup and trial structure. Mice translated a bar randomly presented at a consistent location to the left or right of center of the screen by orienting a wheel with their forepaws (top). Trial epochs defined by the behavioral task events (bottom). B) Wheel velocity across several consecutive example trials. Asterisks indicate onset of stimulus in each trial. Upward and downward deflections indicate leftward and rightward movement, respectively. C) Gray, wheel velocity for 100 randomly-selected trials. Solid and dashed black lines show average leftward and rightward wheel velocity, respectively. D) Fraction of left choices for trials rewarded for leftward and rightward movements (131 sessions from 3 mice). Symbols correspond to mice. Mouse medians are shown in black. E) Distribution of reaction times (defined as duration from stimulus onset to movement initiation) for each mouse. Mean across mice is shown in black. F) Median reaction times as a function of outcome on the current and previous trials for the same sessions plotted in D and E. Grand median for each mouse is shown in black. Symbols correspond to mice as in D. Shorter reaction times correlated with errors on the current trial and correct choices on the previous trial.

Indeed, we found that reaction times were shorter on incorrect compared to correct trials in 2/3 mice (Fig. 1F, left; n=138 sessions; logistic regression, Wald test, *p*<0.05), perhaps reflecting the fact that shorter reaction times did not allow for sufficiently accurate sensorimotor processing. These data were consistent with those collected during a similar head-fixed sensorimotor task by the International Brain Laboratory (International Brain Laboratory et al., 2024) (IBL task: 80%: 67/84 experimental sessions; logistic regression, Wald test, *p*<0.05). Furthermore, given that choices on similar tasks are often influenced by recent stimuli, choice, and immediate trial history (i.e. correct versus incorrect) (Findling et al., 2024; Gold et al., 2008; Thompson et al., 2016), we next asked whether reaction times also reflected such priors in this behavioral task. Indeed, we found that reaction time tended to be longer on trials that followed an unrewarded outcome in 3/3 mice (Fig. 1F, right: n=138 sessions; IBL: 57%: 48/84 experimental sessions logistic regression, Wald test, *p*<0.05). These results indicate that behavior was influenced by prior outcomes in a manner by which mice waited longer to generate their movement choices, specifically on trials that followed unrewarded previous choices.

We then examined task-related neural activity, focusing first on the dSC. We analyzed activity in three task epochs: the baseline (a ≥500 msec period of wheel stillness prior to stimulus onset), choice (from stimulus onset to movement initiation) and execution (from movement initiation to wheel cessation) (Fig. 1; see Methods). Most prominently, we found that the activity of most units differed between the movement and baseline epochs (85%: 98/115 units; Wilcoxon signed-rank test; adjusted p-values (see Methods); Fig. 2A), consistent with dSC recordings during the IBL task (86%: 1118/1303 units; Wilcoxon signed-rank test; adjusted p-values; see Methods). Of the movement-related dSC units, most exhibited a movement-related increase in activity (68%: 67/98 units; IBL: 66%: 739/1118 units), while some exhibited a decrease (32%: 31/98 units; IBL: 34%: 379/1118 units).

**Figure 2.**
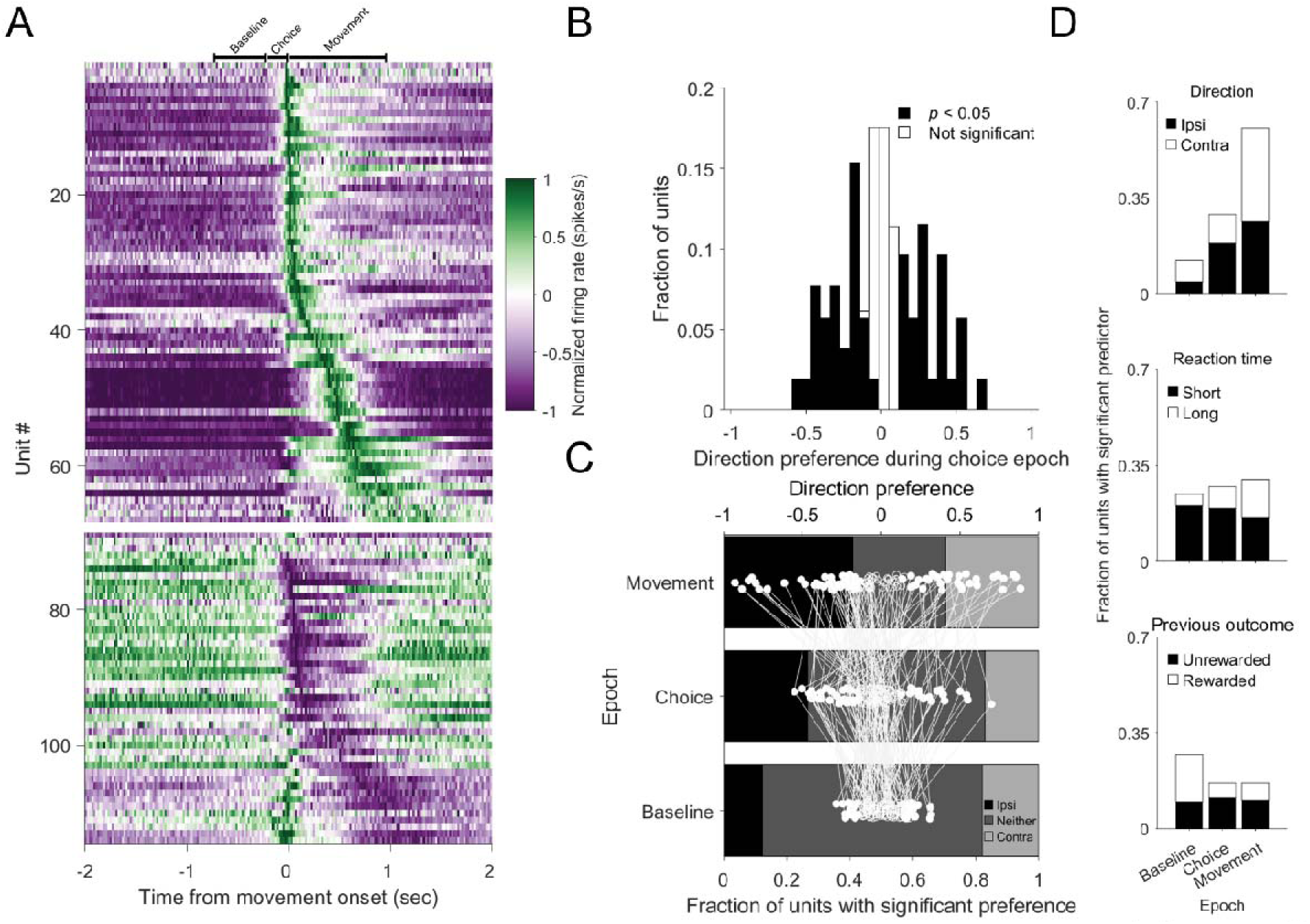
Activity in deep superior colliculus reflect movement direction throughout the trial. A) Normalized activity in 20 msec bins of all dSC units aligned to movement onset. Units exhibiting increasing (upper) and decreasing (lower) responses were separately sorted according to the timing of their response. Bars above heatmap indicate grand median epoch durations. B) Direction preference during the choice epoch. Units with a significant preference are displayed in black. C) Shading shows fraction of dSC units with a significant preference during baseline, choice and movement epochs (lower x-axis). White dots connected by lines show direction preference for each unit in each epoch (upper x-axis). Filled circles indicate significant preference during that epoch. D) Fraction of units in which linear regression analysis of epoch-based firing rates assigned significant weights to movement direction (ipsilateral vs. contralateral; upper), reaction time (middle), and the outcome of the previous trial (rewarded or unrewarded; lower).

We next sought to determine whether and when dSC activity reflected behavioral variables. Given that the dSC is critical for spatial decision making (Basso & May, 2017; Essig et al., 2021; Hoy & Farrow, 2025; Krauzlis et al., 2004; Stubblefield et al., 2013), we first examined whether neural activity exhibited a preference for the direction of the upcoming movement (ipsilateral vs. contralateral, Methods) during the choice epoch. We found that separate subpopulations preferred ipsilateral (preference < 0) and contralateral movement (preference > 0) (Fig. 2B). In addition, many units exhibited direction preference across both the baseline and movement epochs (Fig. 2C), consistent with units in the IBL data set (Baseline: 50%: contralateral: 22%, ipsilateral: 28%; Choice: 52%: contralateral: 19%, ipsilateral: 33%; Movement: 79%: contralateral: 32%; ipsilateral: 47%; n=676 units). Direction preference was more frequent during the choice and movement epochs than the baseline epoch (*p*<0.0001; one-tailed *Z*-test), as expected given that movement direction should not be selected prior to stimulus presentation. Across epochs, units were about as likely to prefer ipsilateral as contralateral movement.

We next examined the relationship between dSC activity in each epoch and several inter-related, task-relevant variables using linear regression models (see Methods; Fig. 2D). Consistent with our direction preference results, we found that the activity of many dSC units was significantly influenced by movement direction, and increasingly so from the baseline to the choice to the movement epochs (Fig. 2D, top; IBL: Baseline: 26%; Choice: 42%; Movement: 63%; n=1303 units). We also found that activity across epochs depended on reaction time (Fig. 2D, middle; IBL: Baseline: 44%; Choice: 47%; Movement: 52%; n=1303 units) and, to a lesser extent, the outcome of the previous trial (Fig. 2D, bottom; IBL: Baseline: 35%; Choice: 27%; Movement: 25%; n=1303 units), reflecting the influence of decision-related and kinematic variables, respectively. Together, these results show that the activity of dSC units reflected several task-related variables throughout the trial, consistent with previous recordings during other orienting tasks (Basso & Wurtz, 1998; Crapse et al., 2018; Felsen & Mainen, 2008; Ikeda & Hikosaka, 2003).

When we performed the same analyses on the population of recorded SNr units (Fig. 3), many results resembled those of the dSC. Activity frequently differed between the movement and baseline epochs (Fig. 3A; 87%: 120/138 units; IBL: 80%: 125/157 units; Wilcoxon signed-rank test; adjusted p-values; see Methods), with most movement-related SNr units exhibiting an increase in activity (88%: 105/120 units; IBL: 73%: 91/125 units) and others exhibiting a decrease (12%: 15/120 units; IBL: 27%: 34/125 units). We also found that the activity across epochs of many SNr units, like dSC units, frequently depended on movement direction (Fig. 3B-D, top; IBL: Baseline: 47%: contralateral: 26%, ipsilateral: 21%; Choice: 51%: contralateral: 29%, ipsilateral: 22%; Movement: 59%: contralateral: 32%, ipsilateral: 26%; n=157 units) and reaction time (Fig. 3D, middle; IBL: Baseline: 38%; Choice: 43%; Movement: 36%; n=157 units).

**Figure 3.**
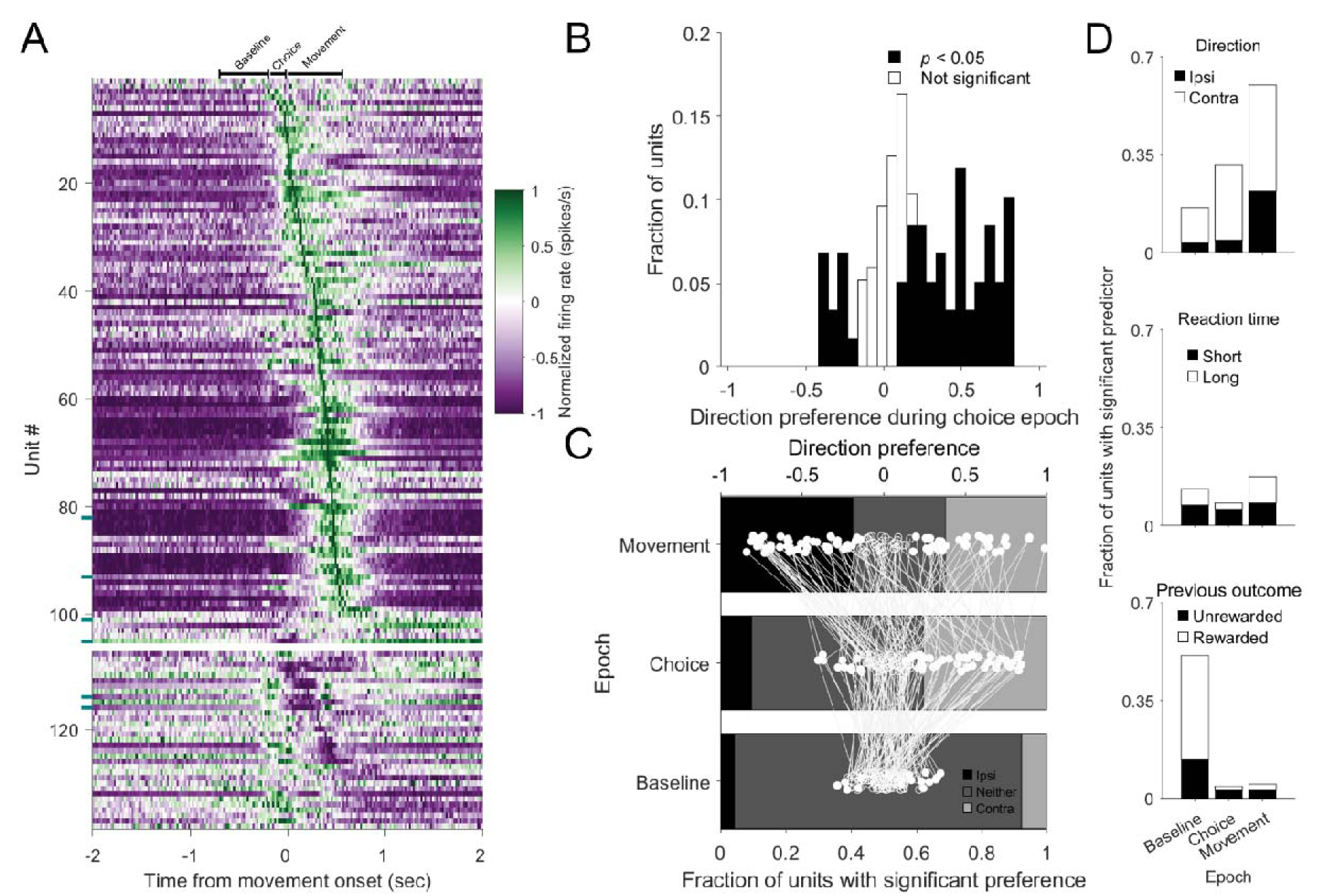
Activity in substantia nigra p*ars reticulata* reflects movement direction and previous trial outcome. A) As in Figure 2, for SNr units. Teal tick marks (left side) indicate SNr units that were optogenetically tagged and thus, putatively project to the ipsilateral dSC. B-D) As in Figure 2, for SNr units.

However, in contrast with our findings in dSC, SNr activity during the baseline epoch frequently depended on the outcome of the previous trial (Fig. 3D, bottom; 51%; 70/137 units; IBL: 29%: 45/157 units). Of these units, 73% (51/70) exhibited increased activity following a rewarded trial, while 27% (19/70) exhibited decreased activity. Such representations are consistent with behavioral strategies sensitive to prior rewards, as observed in this and similar tasks (Findling et al., 2024; Gold et al., 2008; Thompson et al., 2016).

The SNr is thought to influence movements via its inhibitory projection to the dSC, suggesting that SNr and dSC activity would exhibit opposing relationships to movements. For example, the classical disinhibition model holds that phasic release from SNr inhibition permits the increase in dSC activity that initiates movement (Hikosaka & Wurtz, 1983). However, overall, we found that the relationship between neural activity and movement was similar across neurons in the SNr and dSC (Figs. 2 and 3). We therefore examined the role of the SNr-dSC projection in the movements required by our task in two ways.

First, we computed correlations between simultaneously recorded pairs of SNr and dSC units around the time of movement. If dSC is strongly inhibited by SNr during movement, we would expect to frequently observe SNr and dSC pairs with anticorrelated activity. However, after accounting for the correlations exhibited by activity in both regions with movement itself, we found that no pairs of SNr and dSC units were anticorrelated during the movement epoch (all |*r|*<0.2; Pearson correlation; n=2,724 pairs; Fig. 4A,C, right). In contrast, we identified pairs of dSC units that exhibited positively or negatively correlated activity during the movement epoch (Fig. 4B), demonstrating that this analysis is capable of uncovering functional connectivity. Similarly, we found no anticorrelated pairs during the baseline epoch, when we might expect SNr to transmit information about previous outcome to dSC (Fig. 4C, left). Next, in a subset of experiments, we optogenetically tagged putatively dSC-projecting SNr units by expressing ChR2 in SNr neurons and identifying units that exhibited antidromically-elicited spikes in response to dSC light delivery (Methods). Relatively few units were tagged and thus identified as dSC-projecting (n=6/66), and these units exhibited similar task-related activity as the overall population of SNr units (Fig. 3A; blue ticks, see left side of plots). Together these findings are consistent with an inhibitory role of the SNr in contributing to the modulation of dSC activity underlying the movements required by this task, rather than being the primary source of dSC suppression.

**Figure 4.**
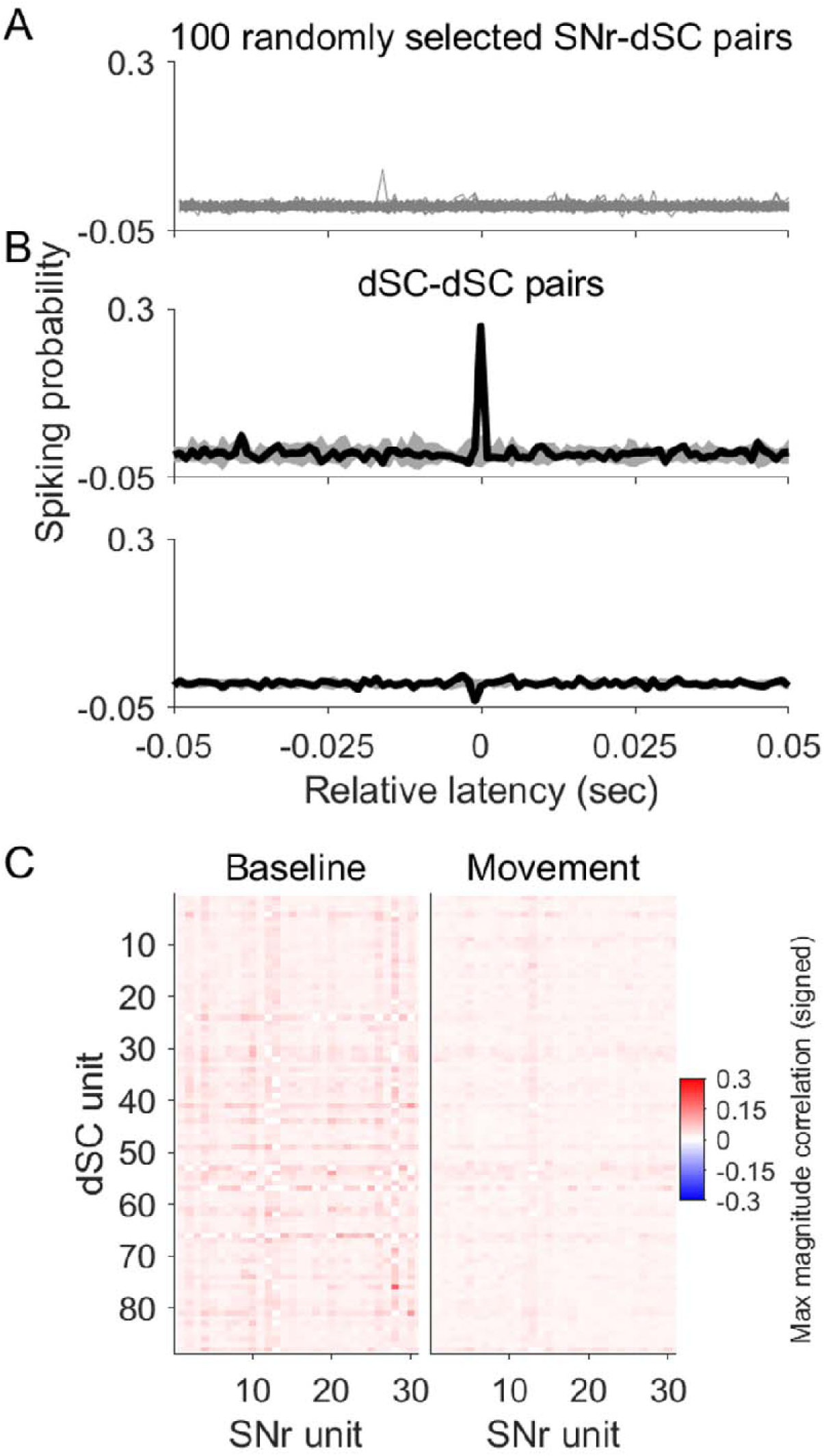
Substantia nigra *pars reticulata* provides limited functional inhibition to deep superior colliculus during head-fixed behavior. A) Cross-correlograms of 100 randomly selected SNr-dSC unit pairs. Probability of a dSC spike is shown as a function of lag from an SNr spike, calculated during the movement epoch. B) Pairs of dSC units exhibited functional excitatory (top) and inhibitory (bottom) coupling during the same (movement) epoch. C) Heat map showing the maximum magnitude of correlation within lags of −50 to 0 msec for each SNr-dSC pair during the baseline (left) and movement (right) epochs.

Finally, to further examine how neural activity depends on task contingencies, we recorded neural activity in additional mice performing one of two variants of the behavioral task. First, based on previous findings that SNr and dSC activity relates differently to movements guided by sensory and non-sensory stimuli (Handel & Glimcher, 2000; Lintz et al., 2019; Lintz & Felsen, 2016), we modified the behavioral task such that the side of stimulus presentation and rewarded movement direction were fixed for several consecutive trials (Methods), increasing the predictability of the rewarded movement direction. We found that mice performed this blocked variant well, frequently selecting the correct, rewarded direction of movement (Fig. 5A) and initiating movement as soon as possible to maximize reward rate. During the movement epoch, we found that dSC activity was frequently modulated – increased or decreased relative to baseline (Fig. 5B) – similar to the original task. We again found that movement-related activity was similar in putatively dSC-projecting SNr units (12/40 were optogenetically tagged in a subset of sessions, Fig 5D) compared to the untagged population (Fig. 5C), although tagged units never exhibited a significant movement-related decrease in activity (n=12), in contrast to untagged units (increasers: 21 units; decreasers: 7 units; Wilcoxon signed-rank test; *p*<0.05). We did not examine activity in the baseline or choice epoch since, by design, mice frequently initiated movement before stimulus presentation.

**Figure 5.**
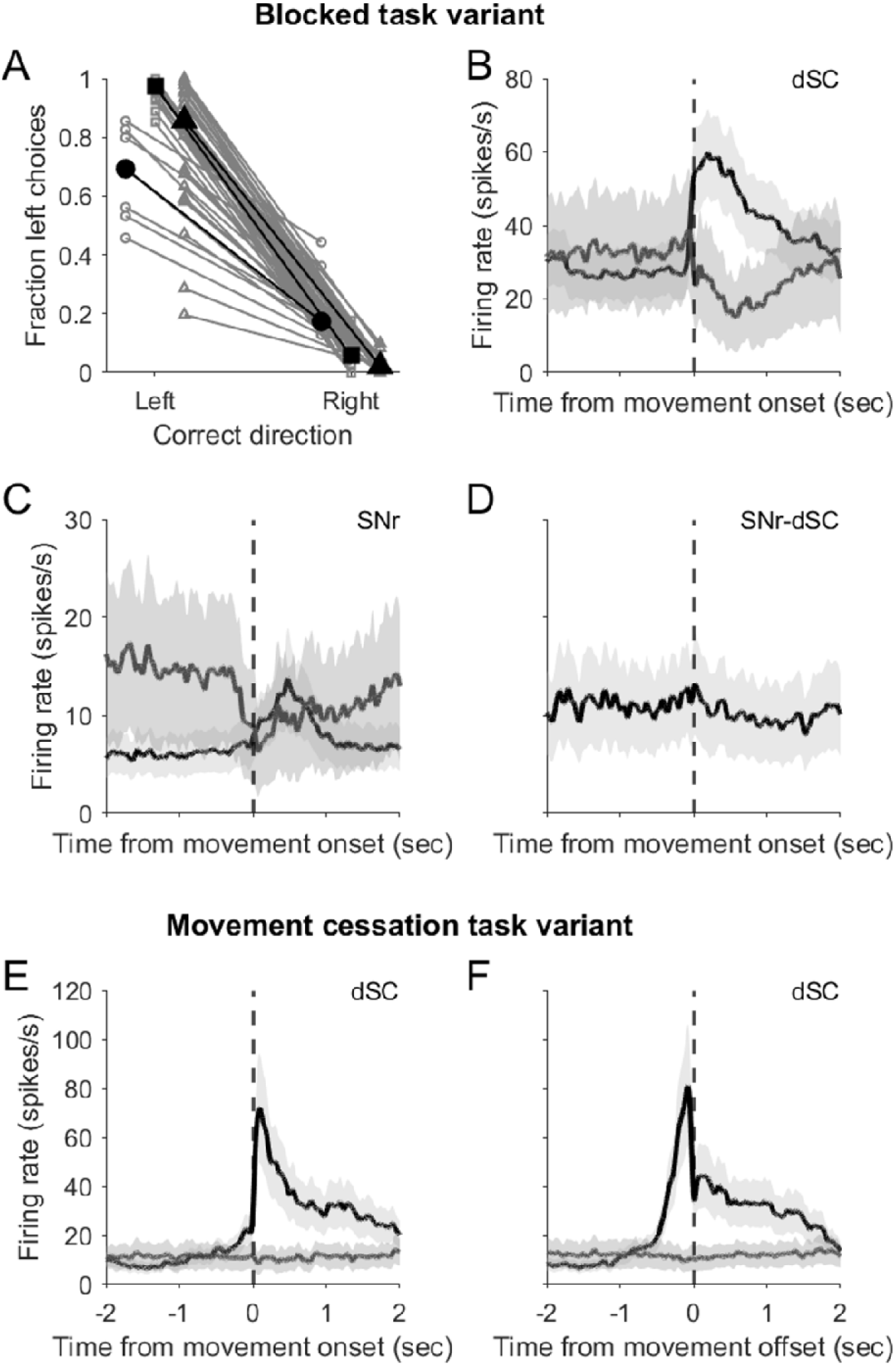
Movement-related activity in substantia nigra *pars reticulata* and deep superio colliculus in variants of the head-fixed task. A) Fraction of left choices for trials rewarded for leftward and rightward movements during the blocked task variant (56 sessions from 3 mice). Symbols correspond to each mouse. Mouse medians are shown in black. B) Population peri-event time histogram (PETH) for dSC units with significant movement-related neural activity, separately for increasers (black, n = 47) and decreasers (dark gray, n = 8). Light gray, ±SEM. C) Same as B but for SNr units not identified as projecting to dSC (n = 21 increasers, black; n = 7 decreasers, dark gray). Light gray, ±SEM. D) PETH for SNr units identified as projecting to dSC (n = 12 increasers, black; 0 decreasers). Light gray, ±SEM. E, F) Population PETH for 9 dSC units with significant movement-onset-(E) and offset-(F) related neural activity (n=6 increasers, black; n=3 decreasers, dark gray), recorded during a task variant requiring movement cessation on target. Light gray, ±SEM.

In the second task variant, we examined whether behavior and neural activity would differ if mice were required to stop their movement on target, rather than overshoot the target as they frequently did in the original task and in the IBL task. Mice were trained accordingly and became adept at stopping their movement when the bar reached the target, and we again found that dSC units were active during the movement (Fig. 5E; SNr units were not recorded during this task). In addition, this variant revealed that dSC activity reflected visually-guided movement endpoint, ramping up towards the end of the movement and sharply decreasing just before the target was acquired (Fig. 5F). These results demonstrate that dSC activity correlates with movement cessation, consistent with the dSC representing distance to the movement target and in stopping movement on target (Choi & Guitton, 2009; Essig & Felsen, 2021; Van Opstal & John, 2023). Together, these data suggest that dSC plays a general role in the goal-directed, orienting forelimb movements common across task variants.

## DISCUSSION

We examined dSC and SNr activity in several variants of a sensorimotor task requiring head-fixed mice to perform orienting forelimb movements. Across task variants and in both brain regions, we found that activity reflected task-relevant variables during the baseline, choice, and movement epochs of the task (Figs 2, 3, 5). These data are consistent with a body of work implicating the involvement of both regions in a broad range of orienting effector movements (Basso & Sommer, 2011; Deniau et al., 2007; Gandhi & Katnani, 2011; Ito et al., 2025; Lintz & Felsen, 2016; Philipp & Hoffmann, 2014; Thomas et al., 2023).

We found that dSC and SNr each contained separate populations of neurons that exhibited either an increase or decrease in activity during movement (Figs 2A, 3A, 5B-F), consistent with previous findings (Handel & Glimcher, 1999; Lintz et al., 2019; Lintz & Felsen, 2016). This observation is also consistent with the inhibitory influence of SNr on downstream targets: for example, SNr neurons with a movement-related increase in activity could provide inhibition to dSC neurons with a movement-related decrease in activity. However, when we examined functional connectivity directly, we did not observe negative correlations between any pairs of SNr and dSC units (Fig. 4A, D). In addition, even our subpopulation of putatively dSC-projecting SNr units did not exhibit activity consistent with the disinhibition model: notably, more SNr-dSC units exhibited movement-related increases in activity, rather than decreases. While it is difficult to rule out alternatives, including that we may not have optimally targeted the dorsolateral region of the SNr that projects most strongly to dSC (Deniau & Chevalier, 1992; Rossi et al., 2016), our data are consistent with a circuit model whereby SNr inhibition modulates, rather than suppresses, dSC activity, at least for the head-fixed, orienting forelimb movements examined here (Basso & Sommer, 2011; Park et al., 2020; Villalobos & Basso, 2022; Yttri & Dudman, 2016, 2018). Within this framework, SNr is one of several brain regions providing task-relevant input to dSC (Basso et al., 2021; Cang et al., 2024; Huda et al., 2020; Wolf et al., 2015).

The modulation of dSC activity by the SNr may serve different functions across phases of the task. During the baseline epoch, the SNr may set the tone of activity in downstream motor regions to bias choices and movements towards the targets most likely to be rewarded (Hikosaka et al., 2006; Lo & Wang, 2006) based on previous choices and outcomes (Fig. 3D). Specifically, reaction times are longer following incorrect trials (Fig. 1F) demonstrating that behavior is influenced by recent history, and previous outcome is also reflected in SNr activity (Fig. 3D), consistent with findings in the IBL task (Findling et al., 2024). Indeed, the subset of dSC units that exhibited direction preference during the baseline epoch (Fig. 2C, D) may reflect this history-dependent bias. During the choice epoch, inhibition from SNr may be one of many inputs influencing which direction of movement is selected, rather than serving as the primary determinant of dSC activity by phasically releasing dSC from tonic inhibition (Wolf et al., 2015). Finally, during the movement, SNr may modulate movement kinematics, such as speed and vigor (Yttri & Dudman, 2018).

Our results support the theory that, even for simple sensorimotor tasks, the relevant neural activity is distributed across subsets of neurons in many brain regions (International Brain Laboratory et al., 2024; Steinmetz et al., 2019). While the activity of the majority of dSC and SNr units was modulated during the task, the activity of many units in each region did not depend on any of the task variables that we examined (Figs. 2D, 3D). These results point to the importance of identifying how the activity of inter-areal neuronal populations, rather than individual neurons, contribute to behavior (Saxena & Cunningham, 2019; Semedo et al., 2020).

We observed some differences from previous studies examining the neural bases of orienting tasks. Most notably, across tasks and across species, previous studies have found that dSC activity reflects contralateral choices (Felsen & Mainen, 2012; Horwitz & Newsome, 2001; Krauzlis et al., 2004; Lintz et al., 2019; Thomas et al., 2023). In contrast, we observed that dSC units were no more likely to exhibit a preference for contralateral than ipsilateral movements during the decision epoch, in our task (Fig. 2B-D) and in the IBL data set. This difference may reflect the fact that the motor plan required for head-fixed mice to turn a wheel with their forelimbs differs considerably from other orienting motor plans; e.g., for directional head movements and saccades. In particular, in a head-fixed context, a contralateral forelimb movement may engage circuitry for (attempting to) orient the head ipsilateral to the limbs (Huda et al., 2020). A contralateral choice may thus require the dSC to coordinate a contralateral forelimb movement with an ipsilateral head movement, which could account for the absence of a contralateral preference across our dSC population. In addition, the forelimb movement required by our task may be represented similarly to reaching movements, which do not elicit strong direction preference (Chaterji et al., 2025; Stuphorn et al., 1999). Nonetheless, these results inform the interpretation of directional tuning observed in head-fixed tasks in mice, as well as comparisons with studies in other species examining the neural bases of orienting movements. Overall, we observed that dSC and SNr activity related to the selection, initiation and execution of the directional, orienting forelimb movements required by several task variants. These results are broadly consistent with studies that examined neural activity during orienting, across several species, and across various tasks and forms of movement. Together, this body of work suggests a general role for modulatory input from SNr to the dSC in mediating orienting movements.

## ACKNOWLEDGMENTS

We thank members of the Felsen lab for constructive comments on the manuscript. We also thank Gabe J. Murphy for insightful conversations and advice on optimizing our behavioral tasks.

## Author contributions

TKD analyzed the data, generated the figures, and drafted the manuscript. TY collected a subset of the data. AB and SH preprocessed subsets of the data. JTD conceived, implemented and supervised the experiments. GF conceived and designed the experiments and analysis, and wrote the manuscript. EAS conceived, implemented and conducted the experiments, collected and analyzed the data, and edited the manuscript.

